# Dentate Gyrus Activin Signaling Mediates the Antidepressant Treatment Response

**DOI:** 10.1101/491613

**Authors:** Mark M. Gergues, Christine N. Yohn, Marjorie R. Levinstein, Benjamin A. Samuels

## Abstract

Antidepressants that target monoaminergic systems, such as selective serotonin reuptake inhibitors (SSRIs), are widely used to treat neuropsychiatric disorders including major depressive disorder, several different anxiety disorders, and obsessive-compulsive disorder. However, these treatments are not ideal because only a subset of patients achieve remission. The reasons why some individuals remit to antidepressant treatments while others do not are unknown. Here, we developed a paradigm to assess antidepressant treatment resistance in mice. Treatment of mice with either chronic corticosterone or chronic social defeat stress effectively induces increased negative valence behaviors. Subsequent chronic treatment with the SSRI fluoxetine reverses these behavioral changes in some, but not all, of the mice, permitting stratification into persistent responders and non-responders to fluoxetine. We found several significant differences in expression of Activin signaling-related genes between responders and non-responders to fluoxetine in the dentate gyrus, a region that we recently reported is critical for the beneficial behavioral effects of fluoxetine. Furthermore, enhancement of Activin signaling in the dentate gyrus converted behavioral non-responders into responders to fluoxetine treatment more effectively than commonly used adjunctive antidepressant treatments, while inhibition of Activin signaling in the dentate gyrus converted responders into non-responders. Taken together, these results demonstrate that the behavioral response to FLX can be bidirectionally modified via targeted manipulations of the dentate gyrus and suggest that molecular- and neural circuit-based modulations of dentate gyrus may provide a new therapeutic avenue for more effective antidepressant treatments or adjunctive therapies.

## INTRODUCTION

Approximately 32-35 million adults in the US population (16%) experience an episode of major depression in their lifetime^1^, and commonly used treatments, such as selective serotonin reuptake inhibitors (SSRIs), are not ideal since only a subset of patients (~33%) achieves remission with initial treatment^2,3^. However, despite this large population of non-remitters, the reasons why some individuals remit to antidepressant treatments while others do not remain unknown. Given that SSRIs are widely used to treat not only major depressive disorder, but also several anxiety disorders and obsessive-compulsive disorder, improving our understanding of the basis of this treatment resistance is of paramount importance. One approach is to decipher the neural circuitry and molecular mechanisms that underlie antidepressant treatment response and resistance.

Several brain regions, including prefrontal and cingulate cortices, amygdala, thalamus, hypothalamus, nucleus accumbens, and hippocampus are implicated in mood disorders through imaging and postmortem studies^4,5^. Within the hippocampus, several preclinical studies in rodents demonstrate that the dentate gyrus (DG) subfield is an essential component of the neural circuitry mediating the antidepressant response. Serotonin 1A receptors on mature dentate gyrus (DG) granule cells are critical mediators of the negative valence behavioral and the neuroendocrine response to the SSRI fluoxetine^6^. Furthermore, chronic treatment with most antidepressants (including SSRIs) stimulates adult neurogenesis in the dentate gyrus (DG)^7,8^. Chronic SSRI treatment increases proliferation of dividing neural precursor cells and promotes maturation and integration of young adult born granule cells (abGCs) into the DG and ablation or impairment of this neurogenic niche results in the loss of some antidepressant-mediated behaviors^7–12^. Direct peptide infusions of brain-derived neurotrophic factor (BDNF), vascular endothelial growth factor (VEGF), or Activin A, yield an antidepressant-like behavioral response^13–17^. Likewise, targeting entorhinal cortex projections to the DG yields an antidepressant-like behavioral response^18^. Optogenetic and chemogenetic manipulations of ventral DG granule cells demonstrate a role in anxiety-related behaviors and stress resilience^19–21^. Serotonin 1B receptors on cholecystokinin (CCK) inhibitory interneurons in the DG are also essential for mediating the negative valence behavioral response to SSRI treatment^22^. Finally, humans suffering from major depressive disorder have fewer DG GCs than controls and DG volume is inversely correlated with the number of depressive episodes^23,24^. Taken together, all of these data indicate that the DG is a principal component of the neural circuitry mediating the antidepressant response. Therefore, both molecular and functional manipulations of the DG will ultimately be important for determining the differences between remitters and non-remitters to antidepressant treatment and may prove instructive for development of new augmentation therapies.

Exposure of rodents to chronic stressful experiences can induce a long-lasting affective state in which there are increases in negative valence behaviors. This negative affective state is often associated with or described as an experimental system to study mood disorders. Several highly distinct stress paradigms are commonly used for this purpose, including chronic mild stress, chronic social defeat stress, and chronic administration of glucocorticoids^11,25–37^. Importantly, these stressed rodents can be treated with antidepressants to reverse the negative valence behaviors and better understand the neural effects of antidepressants. Interestingly, we have noticed that in the Novelty Suppressed Feeding (NSF) behavioral task, SSRI treatment only reverses the effects of chronic glucocorticoid administration in a subset of mice, suggesting that there may be responders and non-responders to antidepressant treatment^29,38^. Therefore, we sought to better understand and characterize this potential treatment resistance phenotype and then to assess differences in the DG between responders and non-responders in order to determine how to manipulate the DG to modify the response to antidepressant treatment.

## RESULTS

### Behavioral Responders and Non-Responders to FLX treatment following CORT administration

To better understand the potential treatment resistance phenotype following chronic stress and antidepressant treatment, we began by exposing a cohort (n=70) of group housed 8-week-old male C57BL/6J mice to chronic administration with either vehicle or corticosterone (CORT, 5mg/kg/day via drinking water). Chronic CORT administration at this dosage induces several negative valence behaviors, including increased latency to feed in NSF and decreased open arm entries and duration in the elevated plus maze (EPM)^11^. We administered vehicle or CORT for 4 weeks and then added either vehicle or the SSRI fluoxetine (FLX, Prozac, 18mg/kg/day) to the treatment paradigm for an additional 3 weeks (timeline of treatments in Figure 1a). As expected, chronic CORT induced an increased latency to feed in NSF relative to vehicle only treated mice (CORT+VEH vs VEH p = 0.004, logrank Mantel-Cox test with Bonferroni correction) and coadministration of CORT and FLX significantly reduced latency to feed relative to CORT treated mice indicative of an antidepressant response (CORT+FLX vs CORT+VEH p < 0.0001, logrank Mantel-Cox with Bonferroni correction) (Figure 1b left). However, closer inspection of the individual latencies of CORT+FLX mice demonstrated a bimodal distribution (Figure 1b right), providing a potential basis for dividing mice into responders and non-responders to FLX treatment groups. Importantly, all mice that received FLX showed similar levels of FLX in their serum (Supplemental Figure 1). Furthermore, food consumption in the home cage was similar among all mice (data not shown).

**Figure 1.**
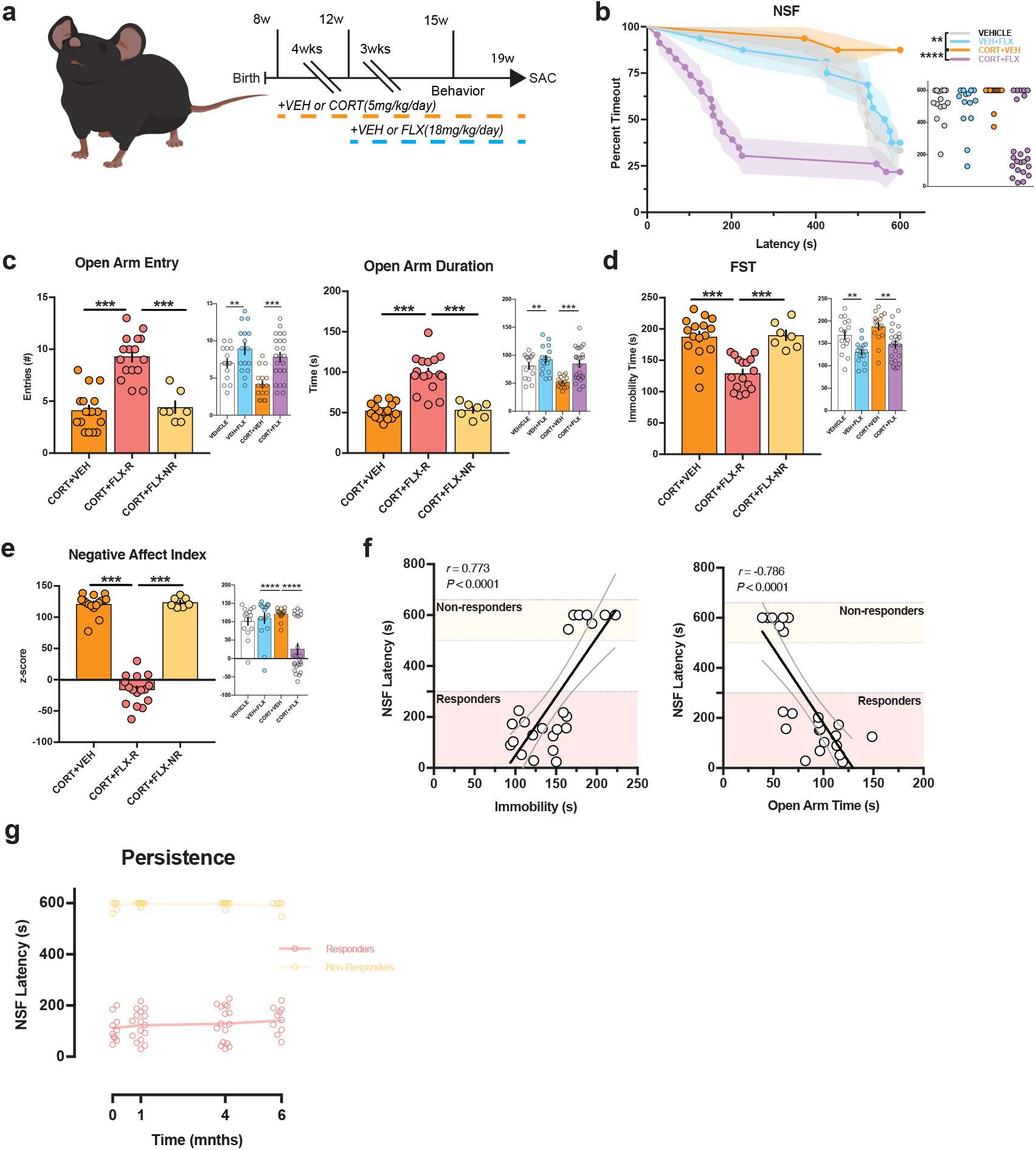
Behavioral Responders and Non-Responders to FLX treatment following CORT administration. (a) Timeline of experiment. (b) Kaplan-Meier survival curve (large panel) and scatterplot (small panel) of NSF data showing individual latency to eat values across all four treatment groups. (c-e) Two-way ANOVA of all treatment groups (small panel) and One-Way ANOVA of CORT+VEH, CORT+FLX responders and CORT+FLX non-responders (large panel) for EPM open arm entries (c left panel), EPM open arm duration (c right panel), FST immobility (d), and Negative Affect Index (e). (f) Regression analyses correlating NSF latency to eat with FST immobility (left) and EPM open arm duration (right). (g) In a separate cohort of CORT+FLX mice, persistence of response was determined by assessing NSF behavior after 3 weeks of FLX (time point 0), and then again 1, 4, and 6 months later. For survival curves, line shading shows SEM of each group (n=15-23 per group). Scatterplots, horizontal lines, and bars show group means with errors bars indicating SEM (n=7-16 per group).

We next exposed the same cohort of C57BL/6J mice to EPM and then the forced swim test (FST), which is a commonly used test of antidepressant efficacy. In the EPM, separate two-way ANOVAs revealed effects of CORT administration and FLX treatment in open arm entries (CORT: *F*_(1,66)_ = 9.69, p = 0.0027, FLX: *F*_(1,66)_ = 19.4, p < 0.0001) and open arm duration (CORT: *F*_(1,66)_ = 10.34, p = 0.002, FLX: *F*_(1,66)_ = 15.42, p = 0.0002) (Figure 1c small panels). To investigate behavioral differences between CORT only treated mice, NSF-defined CORT+FLX responders, and non-responders in the EPM, we next used one-way ANOVAs and found significant differences in open arm entries (*F*_(2,36)_ = 33.24, p < 0.001) and duration (*F*_(2,36)_ = 36.54, *p* < 0.001) (Figure 1c large panels), with Bonferroni-corrected post hoc tests demonstrating that responders had significantly increased open arm entries and duration relative to vehicle treated mice and non-responders (entries and duration: CORT+VEH vs CORT+FLX-R and CORT+FLX-R vs CORT+FLX-NR, all p < 0.001). Non-responders did not show any significant differences relative to vehicle treated mice (entries and duration: CORT+VEH vs CORT+FLX-NR, p > 0.999 for both). These data suggest that FLX response status is conserved across the NSF and EPM. Similarly, in the FST, a two-way ANOVA revealed effects of both CORT administration (*F*_(1,66)_ = 4.83, p = 0.031) and FLX treatment (*F*_(1,66)_ = 22.24, p < 0.0001) (Figure 1d small panel) on immobility over the last four minutes of the six minute test. A separate one-way ANOVA demonstrated significant differences in immobility (*F*_(2,36)_ = 21.02, p < 0.001) (Figure 1d large panel), with Bonferroni-corrected post hoc tests demonstrating that responders had significantly decreased immobility relative to vehicle treated mice and non-responders (CORT+VEH vs CORT+FLX-R and CORT+FLX-R vs CORT+FLX-NR, both p < 0.001). Non-responders did not show any significant differences relative to vehicle treated mice in the FST (CORT+VEH vs CORT+FLX-NR, p > 0.999). Taken together, these data suggest that FLX response status across NSF, EPM, and FST is conserved in CORT-treated mice.

A negative affect index was next used to assess the behavior of this cohort of mice across EPM, NSF, and FST as previously described^39,40^ (Figure 1e). Briefly, z-scores were calculated in each behavioral test (EPM, NSF, FST) by normalizing individual animals against control group averages and standard deviation. Each behavioral test z-score was then averaged for each animal and group averages were calculated. The score shows a more comprehensive analysis of behavior across multiple behavioral modalities where a score above zero represents an animal that shows low open arm entries and time in the EPM, high immobility times in the FST, and longer latency to feed in the NSF task relative to control. A two-way ANOVA revealed significant effects of CORT (*F*_(1,66)_ = 6.41, p = 0.013) and FLX (*F*_(1,66)_ = 12, p = 0.0009) treatment on the negative affect index (Figure 1e small panel). A separate one-way ANOVA demonstrated significant differences in negative affect index (*F*_(2,36)_ = 261.4, p < 0.001), with Bonferroni-corrected post hoc tests demonstrating that responders had a significantly decreased negative affect index relative to vehicle treated mice and non-responders (CORT+VEH vs CORT+FLX-R and CORT+FLX-R vs CORT+FLX-NR, both p < 0.001) (Figure 1e large panel). Non-responders did not have a significantly different negative affect index than vehicle treated mice (CORT+VEH vs CORT+FLX-NR, p > 0.999).

We next directly assessed the relationship between NSF latency to feed and behavioral performance in the EPM and FST among CORT+FLX treated mice. Significant relationships emerged between NSF latency to feed and open arm time (Pearson *r* = −0.786, p < 0.0001, Figure 1f), as well as NSF latency to feed and immobility duration (Pearson *r* = 0.773, p < 0.0001). To characterize these relationships further we ran two separate linear regressions, with NSF latency and open arm time having a linear regression line (*y* = −6.02x + 780, *F*_(1,21)_ = 33.9, p < 0.0001), and FST linear regression line (*y* = 4.63x − 415, *F*_(1,21)_ = 31.1, p < 0.0001) (Figure 1f) .

Finally, we wanted to assess whether the responder and non-responder phenotypes persisted across several months (Figure 1g). To this end, we exposed a new cohort of C57BL/6J mice to CORT+FLX as in Figure 1a, and then assessed behavior in NSF. Similar to the cohort in Figure 1b, the CORT+FLX mice displayed a bimodal distribution of latencies to feed. These mice then remained on CORT+FLX and were retested several times in the NSF. Importantly, the responder vs non-responder behavioral distinction persists for at least 6 months (repeated measures ANOVA reveals significant effect of response status [*F*_(1, 22)_ = 4350, p < 0.0001], but not time [*F*_(3, 66)_ = 0.255, p = 0.8573]). Therefore, the CORT+FLX paradigm permits definition of persistent responders and non-responders to FLX treatment and potentially allows for additional manipulations in attempts to convert non-responders into responders.

### Dentate Gyrus mRNA expression of Activin signaling components correlates with behavioral response to FLX treatment following CORT administration

We previously published a preliminary microarray study assessing DG gene expression in CORT+VEH and CORT+FLX-treated mice^38^. However, when we looked at CORT+FLX responders vs non-responders in these microarray data, pathway analyses suggested that there were differences in multiple components of DG Activin signaling. We were particularly interested in further analyzing this pathway because previous reports have demonstrated that some Activin signaling components are altered in DG by antidepressant treatment and that acute Activin A infusions have antidepressant-like effects in FST^16,17^.

To fully characterize Activin signaling in DG of responders and non-responders, we prepared a new cohort of vehicle, CORT+VEH, and CORT+FLX treated mice (behavioral data in Supplemental Figure 2), and prepared DG RNA following behavioral testing. Two-way ANOVAs revealed significant effects of FLX treatment on DG expression of Activin A (Figure 2a middle panel, *F*_(1,62)_ = 85.51, p < 0.0001), the Activin receptors acvr1a (Figure 2b middle panel, *F*_(1,62)_ = 28.15, p < 0.0001) and acvr1c (Figure 2d middle panel, *F*_(1,62)_ = 35.37, p < 0.0001), and the intracellular signaling protein smad3 (Figure 2f middle panel, *F*_(1,62)_ = 45.19, p < 0.0001) and of CORT administration on Activin A (Figure 2a middle panel, *F*_(1,62)_ = 5.01, p = 0.0288) and acvr1b (Figure 2c middle panel, *F*_(1,62)_ = 7.285, p = 0.009). Separate one-way ANOVAs were next used to compare DG expression of these genes in CORT+FLX responders, CORT+FLX non-responders, and CORT+VEH mice. These analyses revealed significant group differences in Activin A (Figure 2a left panel, *F*_(2,36)_ = 81.68, p < 0.001), acvr1a (Figure 2b left panel, *F*_(2,36)_ = 34.7, p < 0.001), acvr1c (Figure 2d left panel, *F*_(2,36)_ = 56.97, p < 0.001), smad2 (Figure 2e left panel, *F*_(2,36)_ = 23.73, p < 0.001), and smad3 (Figure 2f left panel, *F*_(2,36)_ = 72.33, p < 0.001). Interestingly, CORT+FLX responders showed increased expression of Activin A (Figure 2a), acvr1a (Figure 2b), acvr1c (Figure 2d), and smad3 (Figure 2f) (p < 0.001 for all, Bonferroni corrected) relative to CORT only treated mice and non-responders to CORT+FLX. When comparing CORT+FLX non-responders to CORT+VEH mice, we found a significant difference in activin A (p = 0.047, Bonferroni corrected) and smad2 expression (p < 0.001, Bonferroni corrected), but not in acvr1a, acvr1b, acvr1c, or smad3 expression (all p > 0.999, Bonferroni corrected). Finally, we directly compared NSF latency to feed with DG expression of these genes. Significant relationships emerged between NSF latency to feed and expression of activin A (Pearson *r* = −0.817, p < 0.0001), with linear regression line (*y* = −3.23x + 2480, *F*_(1,23)_ = 46.2, p < 0.0001) (Figure 2a right panel), acvr1a (Pearson *r* = −0.76, p < 0.0001), with linear regression line (*y* = −0.219x + 243, *F*_(1,25)_ = 31.5, p < 0.0001) (Figure 2b right panel), acvr1c (Pearson *r* = −0.815, p < 0.0001), with linear regression line (*y* = −0.271x + 269, *F*_(1,23)_ = 49.9, p < 0.0001) (Figure 2d right panel), smad2 (Pearson *r* = 0.727, p < 0.0001), with linear regression line *(y* = 0.108x + 69.3, *F*_(1,23)_ = 49.9, p < 0.0001) (Figure 2e right panel), and smad3 (Pearson *r* = −0.858, p < 0.0001), with linear regression line (*y* = −0.221x + 240, *F*_(1,23)_ = 49.9, p < 0.0001) (Figure 2f right panel). Taken together, all of these data demonstrate that DG Activin signaling is significantly different between responders and non-responders to FLX treatment.

**Figure 2.**
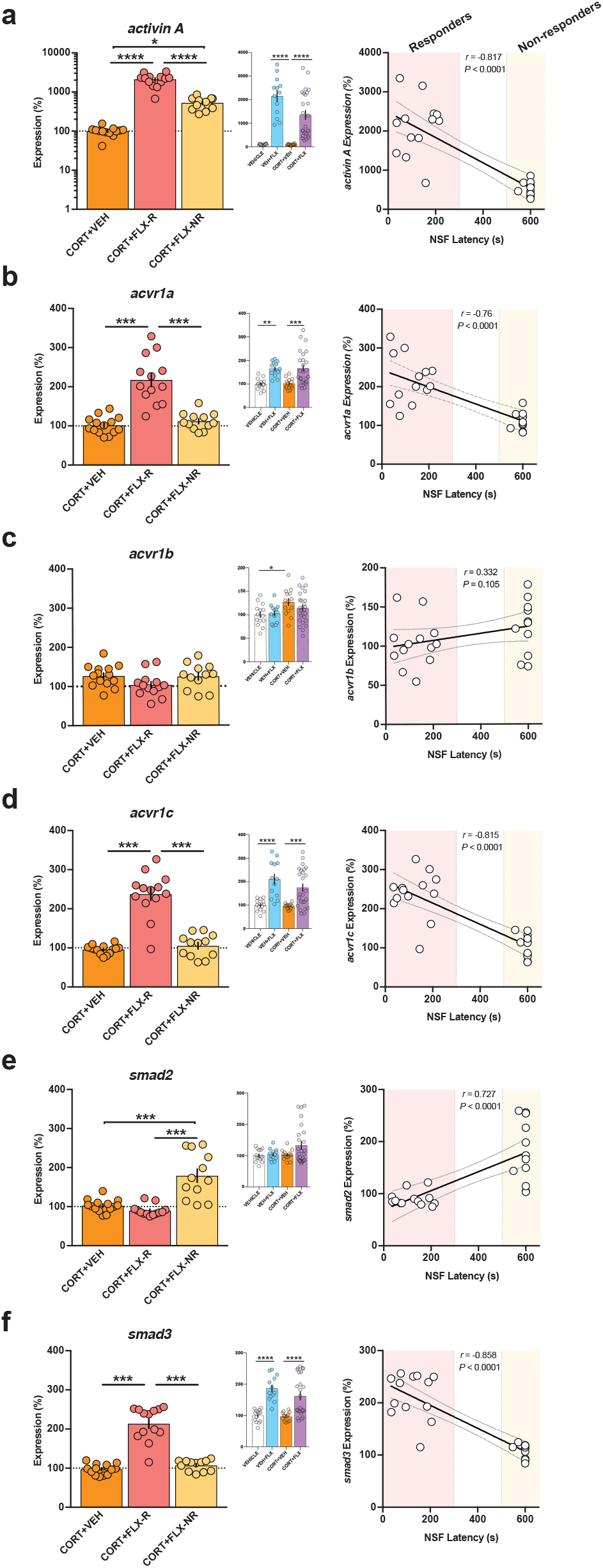
Dentate Gyrus mRNA expression of Activin signaling components correlates with behavioral response to FLX treatment following CORT administration. (a-f) Two-Way ANOVA of all treatment groups (middle panels), One-Way ANOVA of CORT+VEH, CORT+FLX responders and CORT+FLX non-responders (left panels), and regression analyses correlating NSF latency to eat with DG mRNA expression of activin A (a), acvr1a (b), acvr1b (c), acvr1c (d), smad2 (e), and smad3 (f). Scatterplots, horizontal lines, and bars show group means with errors bars indicating SEM (n=12-14 per group).

### Responders and Non-Responders to FLX treatment following chronic social defeat stress replicate the behavioral and DG Activin signaling expression data from the CORT administration paradigm

To confirm that these effects on behavior and Activin signaling were due to differential responses to FLX treatment and not a direct or secondary effect of CORT administration, we next repeated these experiments using a distinct chronic stress paradigm. Chronic social defeat stress (CSDS) is a widely used stress paradigm that involves exposing mice to multiple daily defeats by a conspecific from a larger, more aggressive strain. To this end, we exposed a large cohort (n=125) of 8-week-old male C57BL/6J mice to 10 days of either control or CSDS by CD1 male mice prescreened for aggressive behavior (timeline in Figure 3a). The C57BL/6J mice exposed to CSDS interacted with CD1 aggressors for 5 minutes per day and then were cohoused with the CD1 aggressors separated by a transparent divider for further sensory exposure. Following the 10 days of control or CSDS, the C57BL/6J mice were next exposed to a social interaction test, which indicated that n=35 of the CSDS exposed mice were susceptible (SUS) to the CSDS (Supplemental Figure 3). Control and SUS mice were next administered either VEH or FLX (18mg/kg/day) for 3 weeks and then exposed to a series of negative behavior valence tests. Similar to CORT, SUS mice had an increased latency to feed in NSF relative to control (SUS+VEH vs VEH, p = 0.0003, logrank Mantel-Cox test with Bonferroni correction) (Figure 3b left) and administration of FLX to SUS mice significantly reduced latency to feed (SUS+FLX vs SUS+VEH, p < 0.0001, logrank Mantel-Cox test with Bonferroni correction). Interestingly, similar to CORT, the individual latencies of the SUS+FLX mice showed a bimodal distribution indicative of responders and non-responders to FLX treatment (Figure 3b right).

**Figure 3.**
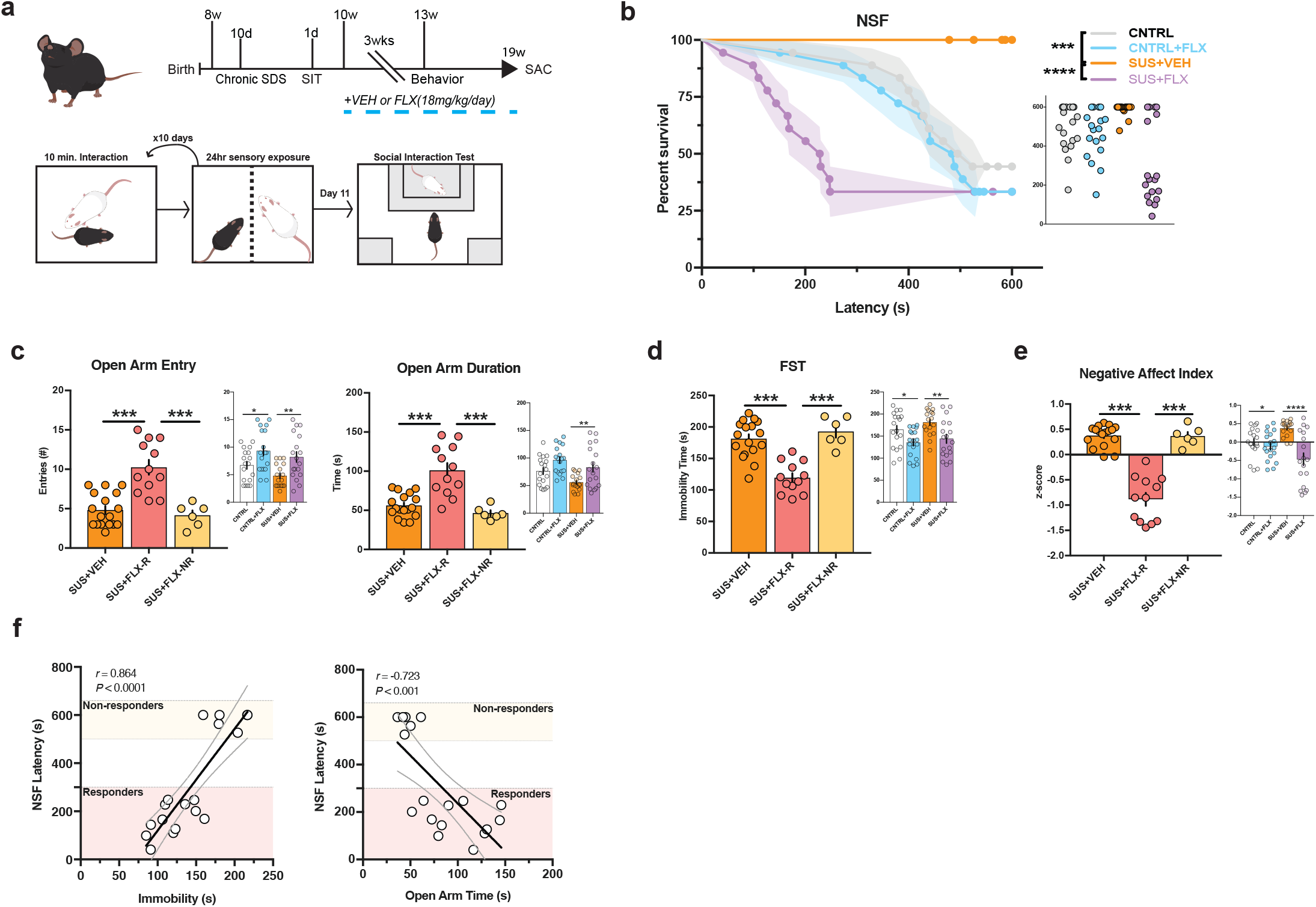
Behavioral Responders and Non-Responders to FLX treatment following. CSDS (a) Timeline of experiment and diagram of CSDS and Social Interaction (SIT) protocol. (b) Kaplan-Meier survival curve (large panel) and scatterplot (small panel) of NSF data showing individual latency to eat values across all four treatment groups. (c-e) Two-way ANOVA of all treatment groups (small panel) and One-Way ANOVA of SUS+VEH, SUS+FLX responders and SUS+FLX non-responders (large panel) for EPM open arm entries (c left panel), EPM open arm duration (c right panel), FST immobility (d), and Negative Affect Index (e). (f) Regression analyses correlating NSF latency to eat with FST immobility (left) and EPM open arm duration (right). For survival curves, line shading shows SEM of each group (n=12-14 per group). Scatterplots, horizontal lines, and bars show group means with errors bars indicating SEM (n=6-18 per group).

The same cohort of mice were next exposed to EPM and FST, and two-way ANOVAs revealed significant effects of FLX treatment for EPM open arm entries (*F*_(1,67)_ = 14.6, p = 0.0003) (Figure 3c small panel), EPM open arm duration (*F*_(1,67)_ = 13.35, p = 0.0005) (Figure 3c small panel), and FST immobility (*F*_(1,67)_ = 14.68, p = 0.00030) (Figure 3d small panel) and of CSDS on EPM open arm duration (*F*_(1,67)_ = 6.993, p = 0.0102) (Figure 3c small panel). Subsequent one-way ANOVAs revealed significant differences in EPM open arm entries (*F*_(2,32)_ = 20.17, p < 0.001) (Figure 3c large panel), EPM open arm duration (*F*_(2,32)_ = 19.12, p < 0.001) (Figure 3c large panel), and FST immobility (*F*_(2,32)_ = 23.34, p < 0.001) (Figure 3d large panel), with SUS+FLX responders showing increased EPM open arm entries and duration and decreased FST immobility relative to SUS+FLX non-responders and SUS+VEH mice (p < 0.001 for all, Bonferroni corrected). SUS+FLX non-responders were not significantly different than SUS+VEH mice in EPM open arm entries, EPM open arm duration, and FST immobility (p > 0.999 for all, Bonferroni corrected). A two-way ANOVA assessing negative affect index in this cohort of mice demonstrated a significant effect of FLX treatment (*F*_(1,67)_ = 18.8, p < 0.0001) (Figure 3e small panel). The subsequent one-way ANOVA found significant group differences (*F*_(2,32)_ = 64.66, p < 0.001) (Figure 3e large panel), with SUS+FLX responders showing a decreased negative affect index relative to SUS+VEH and SUS+FLX nonresponders (p < 0.001 for both, Bonferroni corrected). SUS+FLX non-responders were not significantly different than SUS+VEH mice (p > 0.999, Bonferroni corrected). Significant relationships also emerged when we directly compared NSF latency to feed to EPM open arm duration (Pearson *r* = −0.723, p = 0.0007), with linear regression line (*y* = −4.05x + 641, *F*_(1,16)_ = 17.5, p = 0.0007), and to FST immobility (Pearson *r* = 0.864, p < 0.0001), with linear regression line (*y* = 4.22x − 302, *F*_(1,16)_ = 47.2, p < 0.0001) (Figure 3f). Taken together, these data replicate the CORT behavioral data and demonstrate that FLX response status across NSF, EPM, and FST is conserved in mice susceptible to CSDS.

We next assessed mRNA expression of Activin signaling components in the DG of the CSDS cohort of mice. Two-way ANOVAs revealed significant effects of FLX treatment on DG expression of Activin A (Figure 4a middle panel, *F*_(1,26)_ = 37.45, p < 0.0001), acvr1a (Figure 4b middle panel, *F*_(1,26)_ = 7.717, p = 0.0127), acvr1c (Figure 4d middle panel, *F*_(1,26)_ = 15.58, p = 0.0005), and smad3 (Figure 4f middle panel, *F*_(1,26)_ = 12.64, p = 0.0015) and of CSDS on acvr1b (Figure 4c middle panel, *F*_(1,26)_ = 5.808, p = 0.023). Subsequent one-way ANOVAs found significant group differences for Activin A (Figure 4a left panel, *F*_(2,15)_ = 54.65, p < 0.001), acvr1a (Figure 4b left panel, *F*_(2,15)_ = 9.82, p = 0.002), acvr1b (Figure 4c left panel, *F*_(2,15)_ = 3.78, p = 0.047), acvr1c (Figure 4d left panel, *F*_(2,15)_ = 16.24, p < 0.001), smad2 (Figure 4e left panel, *F*_(2,15)_ = 3.769, p = 0.047), and smad3 (Figure 4f left panel, *F*_(2,15)_ = 23.38, p < 0.001). Interestingly, SUS+FLX responders showed increased expression of Activin A (Figure 4a), acvr1a (Figure 4b, SUS+VEH vs SUS+FLX-R (p = 0.004), SUS+FLX-R VS SUS+FLX-NR (p = 0.006)), acvr1c (Figure 4d), and smad3 (Figure 4f) (all Bonferroni corrected, p < 0.001 for all except acvr1a) relative to SUS+VEH mice and SUS+FLX non-responders. By contrast, similar to CORT, expression of acvr1a, acvr1c, and smad3 were not significantly different between SUS+FLX non-responders and SUS+VEH mice (p > 0.999 for all, Bonferroni corrected). Activin A expression was significantly increased in SUS+FLX non-responders relative to SUS+VEH mice (p = 0.035, Bonferroni corrected).

**Figure 4.**
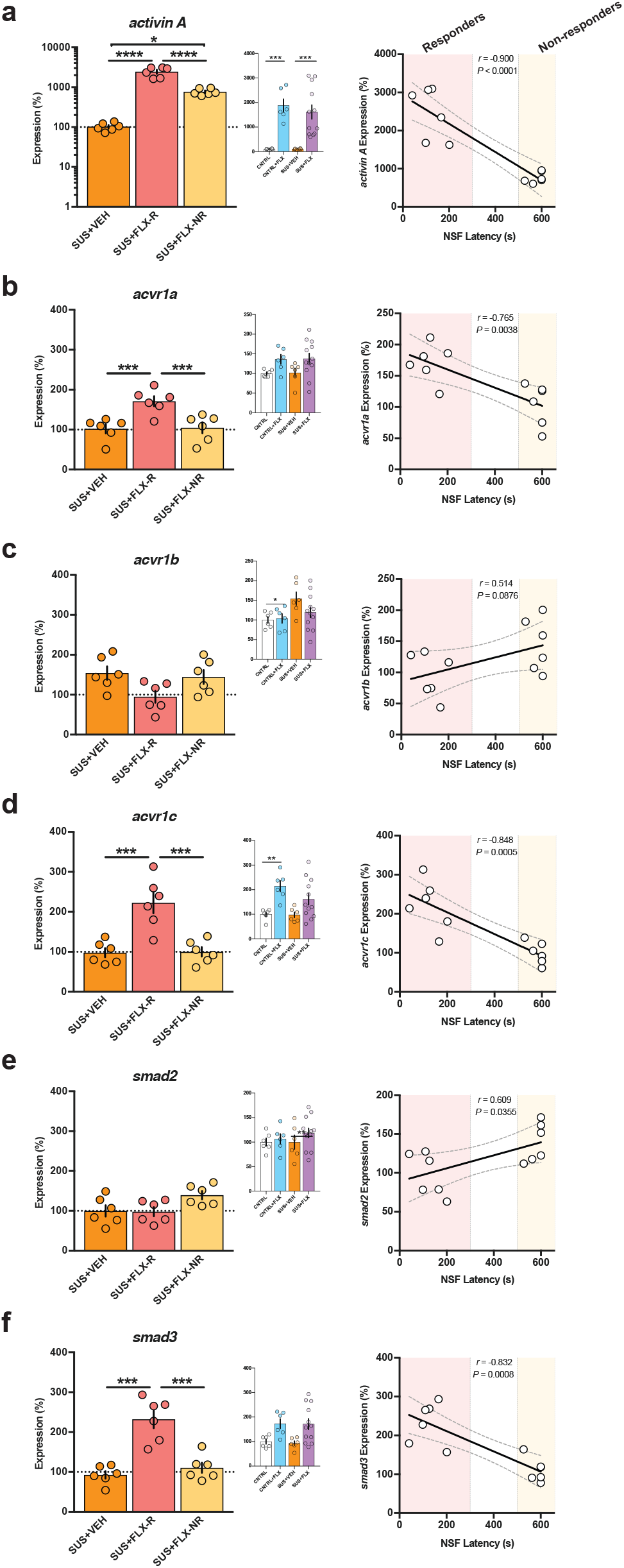
Dentate Gyrus mRNA expression of Activin signaling components correlates with behavioral response to FLX treatment following CSDS administration. (a-f) Two-Way ANOVA of all treatment groups (middle panels), One-Way ANOVA of CORT+VEH, CORT+FLX responders and CORT+FLX non-responders (left panels), and regression analyses correlating NSF latency to eat with DG mRNA expression of activin A (a), acvr1a (b), acvr1b (c), acvr1c (d), smad2 (e), and smad3 (f). Scatterplots, horizontal lines, and bars show group means with errors bars indicating SEM (n=6-12 per group).

Finally, we directly compared NSF latency to feed with DG expression of these genes in responders and non-responders. Significant relationships emerged between NSF latency to feed and expression of activin A (Pearson *r* = −0.900, p < 0.0001), with linear regression line (*y* = −3.70x + 2920, *F*_(1,10)_ = 42.5, p < 0.0001) (Figure 4a right panel), acvr1a (Pearson *r* = −0.765, p = 0.0038), with linear regression line (*y* = −0.145x + 189, *F*_(1,10)_ =14.1, p = 0.0038) (Figure 4b right panel), acvr1c (Pearson *r* = −0.848, p = 0.0005), with linear regression line (*y* = −0.278x + 259, *F*_(1,10_ = 25.7, p = 0.0005) (Figure 4d right panel), and smad3 (Pearson *r* = −0.832, p = 0.0008), with linear regression line (*y* = −0.261x + 263, *F*_(1,10)_ = 22.4, p = 0.0008) (Figure 4f right panel). Taken together, these data replicate the CORT Activin data and demonstrate that DG Activin signaling is significantly different between responders and non-responders to FLX treatment. Furthermore, in two distinct stress paradigms, FLX response status is conserved across behavior and DG Activin signaling.

### Chronic Activin A infusions into DG convert FLX non-responders into responders

Since our gene expression data indicate that several components of DG Activin signaling, including Activin A itself, are decreased in FLX non-responders relative to responders, we wanted to test whether this altered signaling underlies the lack of behavioral response to FLX. Acute Activin A infusions directly into DG yield an antidepressant-like response in FST^16,17^, so we reasoned that development of a chronic Activin A infusion paradigm into DG could potentially convert non-responders to FLX into responders. Since CORT+FLX response status persists for at least six months, we exposed a large cohort of C57BL/6J mice to CORT+FLX, and then non-responders (n=36) received bilateral cannula implants and were infused once daily for two weeks with either vehicle (0.1% BSA), Activin A peptide (1.0 μg per hemisphere) into DG, or Activin A peptide (1.0 μg per hemisphere) into CA1 (timeline in Figure 5a). These mice were then exposed to NSF, EPM, and FST. Remarkably, CORT+FLX non-responders that received chronic Activin A infusions into DG had reduced latency to eat in the NSF relative to non-responders that received vehicle (p < 0.0001, logrank Mantel-Cox test with Bonferroni correction) or chronic Activin A infusions into CA1 (p < 0.0001, logrank Mantel-Cox test with Bonferroni correction) (Figure 5b). Closer inspection of individual latencies demonstrated that all CORT+FLX non-responders that received DG Activin A infusions were converted into responders in the NSF. Group differences were also observed in the EPM for open arm entries (*F*_(2,33)_ = 31.6, p < 0.0001, Figure 5c) and duration (*F*_(2,33)_ = 58.9, p < 0.0001, Figure 5c) and in the FST for immobility (*F*_(2,33)_ = 57.4, p < 0.0001, Figure 5d). CORT+FLX non-responders that received DG Activin A infusions had increased open arm entries and duration and decreased immobility relative to non-responders that received vehicle (p < 0.0001 for all, Bonferroni corrected) or chronic Activin A infusions into CA1 (p < 0.0001 for all, Bonferroni corrected). These data indicate that non-responders to CORT+FLX were converted into responders in NSF, EPM, and FST. The negative affect index also demonstrated group differences (*F*_(2,33)_ = 647, p < 0.0001), with CORT+FLX non-responders that received DG Activin A infusions showing reduced negative affect relative to CORT+FLX non-responders that received vehicle or Activin A infusions into CA1 (p < 0.0001 for both, Bonferroni corrected). Significant relationships also emerged when we directly compared NSF latency to feed to EPM open arm duration (Pearson *r* = −0.873, p < 0.0001), with linear regression line *(y =* −5.81x − 864, *F*_(1,34)_ = 109, p < 0.0001), and to FST immobility (Pearson *r* = 0.891, p < 0.0001), with linear regression line (*y* = 3.77x − 160, *F*_(1,34)_ = 131, p < 0.0001) (Figure 5f). Taken together, these data demonstrate that supplementing Activin signaling in DG can convert FLX non-responders into responders across NSF, EPM, and FST.

**Figure 5.**
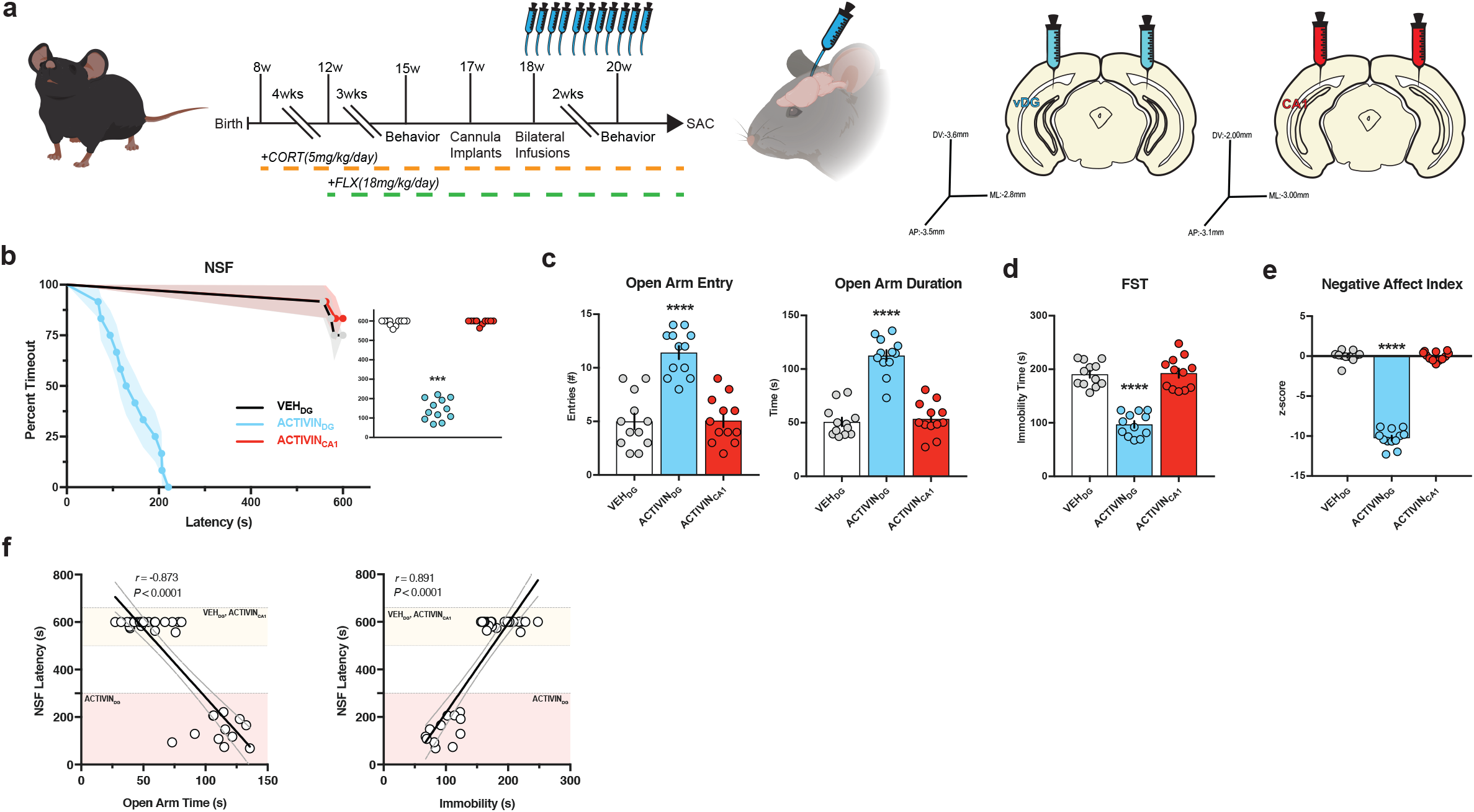
Chronic Activin A infusions into DG convert FLX non-responders into responders. (a) Timeline of experiment and coordinates of infusions for ventral DG and ventral CA1. (b) Kaplan-Meier survival curve (large panel) and scatterplot (small panel) of NSF data showing individual latency to eat values across all three non-responder treatment groups. (c-e) One-Way ANOVA of VEH, Activin A infusions into DG (ACTIVIN_DG_), and Activin A infusions into CA1 (ACTIVIN_CA1_) for EPM open arm entries (c left panel), EPM open arm duration (c right panel), FST immobility (d), and Negative Affect Index (e). (f) Regression analyses correlating NSF latency to eat with EPM open arm duration (left) and FST immobility (right). For survival curves, line shading shows SEM of each group (n=12 per group). Scatterplots, horizontal lines, and bars show group means with errors bars indicating SEM.

### Inhibition of Activin signaling in DG converts FLX responders into non-responders

Since chronic Activin A infusions into DG can convert FLX non-responders into responders, we next wanted to test whether DG Activin signaling was necessary for the behavioral response to FLX. Specifically, we sought to determine whether inhibition of Activin signaling in DG converts FLX responders into non-responders. Inhibin is an endogenously occurring protein complex that has nearly opposite biological effects to Activin^41^. Inhibin binds directly to Activin receptor complexes, where Activin and Inhibin act as mutual antagonists to each other^41^. Therefore, a cohort of C57BL/6J CORT+FLX responders (n=36) received bilateral cannula implants and were infused once daily for two weeks with either vehicle (0.1% BSA), Inhibin A peptide (1.0 μg per hemisphere) into DG, or Inhibin A peptide (1.0 μg per hemisphere) into CA1 (timeline in Figure 6a). These mice were then exposed to NSF, EPM, and FST. Excitingly, CORT+FLX responders that received chronic Inhibin A infusions into DG had increased latency to eat in the NSF relative to non-responders that received vehicle (p < 0.0001, logrank Mantel-Cox test with Bonferroni correction) or chronic Inhibin A infusions into CA1 (p < 0.0001, logrank Mantel-Cox test with Bonferroni correction) (Figure 6b). Group differences were also observed in the EPM for open arm entries (*F*_(2,33)_ = 12.9, p < 0.0001, Figure 6c) and duration (*F*_(2,33)_ = 24.1, p < 0.0001, Figure 6c) and in the FST for immobility (*F*_(2,33)_ = 39.4, p < 0.0001, Figure 6d). CORT+FLX responders that received DG Inhibin A infusions had decreased open arm entries and duration and increased immobility relative to responders that received vehicle (p = 0.0003 for open arm entries, p <0.0001 for open arm duration and immobility, Bonferroni corrected) or chronic Inhibin A infusions into CA1 (p = 0.0003 for open arm entries, p < 0.0001 for open arm duration and immobility, Bonferroni corrected). The negative affect index also demonstrated group differences (*F*_(2,33)_ = 120, p < 0.0001) (Figure 6e), with CORT+FLX responders that received DG Inhibin A infusions showing increased negative affect relative to CORT+FLX responders that received vehicle or Inhibin A infusions into CA1 (p < 0.0001 for both, Bonferroni corrected). Significant relationships also emerged when we directly compared NSF latency to feed to EPM open arm duration (Pearson *r* = −0.776, p < 0.0001), with linear regression line (*y* = −5.99x − 748, *F*_(1,34)_ = 51.3, p < 0.0001), and to FST immobility (Pearson *r* = 0.864, p < 0.0001), with linear regression line (*y* = 4.46x − 391, *F*_(1,34)_ = 100, p < 0.0001) (Figure 6f). Taken together, these data demonstrate that Activin signaling in the DG is necessary for the behavioral effects of FLX treatment as directly inhibiting Activin signaling in DG converts FLX responders into non-responders across NSF, EPM, and FST. Furthermore, these data demonstrate that FLX behavioral response status can be bidirectionally modified by manipulating DG Activin signaling.

**Figure 6.**
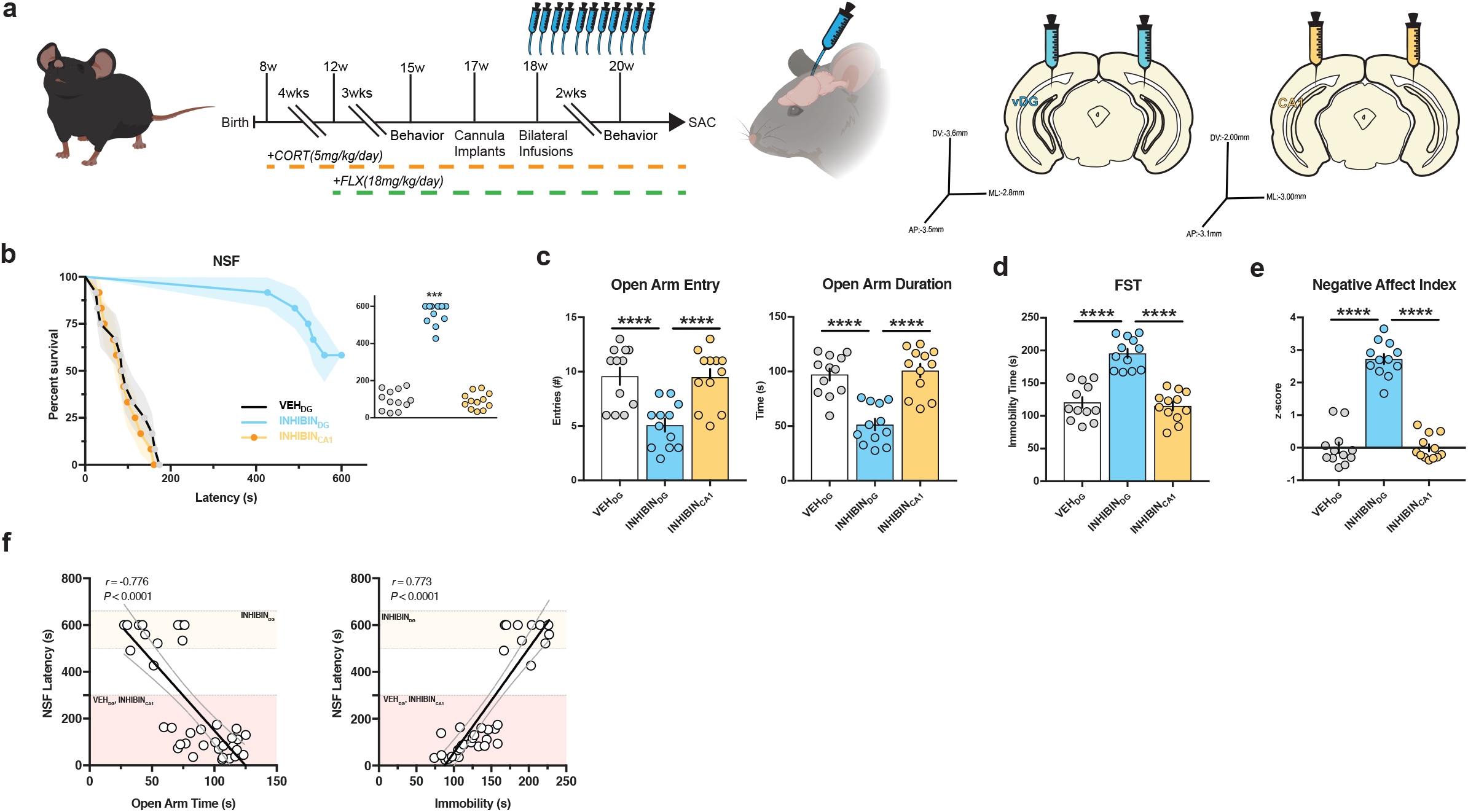
Inhibition of Activin signaling in DG converts FLX responders into non-responders. (a) Timeline of experiment and coordinates of infusions for ventral DG and ventral CA1. (b) Kaplan-Meier survival curve (large panel) and scatterplot (small panel) of NSF data showing individual latency to eat values across all three responder treatment groups. (c-e) One-Way ANOVA of VEH, Inhibin A infusions into DG (INHIBIN_DG_), and Inhibin A infusions into CA1 (INHIBIN_CA1_) for EPM open arm entries (c left panel), EPM open arm duration (c right panel), FST immobility (d), and Negative Affect Index (e). (f) Regression analyses correlating NSF latency to eat with EPM open arm duration (left) and FST immobility (right). For survival curves, line shading shows SEM of each group (n=12 per group). Scatterplots, horizontal lines, and bars show group means with errors bars indicating SEM.

### Coinfusion of Activin A and Inhibin A into Non-Responders blocks the effects of Activin A on behavior

To confirm that the effects of chronic DG Activin A infusions into CORT+FLX nonresponders were directly due to manipulation of Activin signaling as opposed to an off-target effect, we next assessed whether chronic coinfusion of Inhibin A and Activin A blocked the behavioral effects seen with Activin A alone. To this end, a cohort of C57BL/6J CORT+FLX non-responders (n=48) received bilateral cannula implants and were infused once daily for two weeks with either vehicle (0.1% BSA), Activin A peptide into DG (1.0 μg per hemisphere), Inhibin A peptide (1.0 μg per hemisphere) into DG, or both Activin A and Inhibin A peptides (1.0 μg of each per hemisphere) into DG (timeline in Figure 7a). These mice were then exposed to NSF, EPM, and FST. In the NSF, chronic DG Activin A infusions into CORT+FLX nonresponders decreased latency to eat relative to vehicle, Inhibin A, and Activin A+Inhibin infusions (p < 0.0001 for all, logrank Mantel-Cox test with Bonferroni correction) (Figure 7b). Activin A+Inhibin A coinfusions were not significantly different than vehicle (p = 0.129, log-rank Mantel-Cox test with Bonferroni correction) or Inhibin A (p = 0.031, log-rank Mantel-Cox test not significant with Bonferroni correction). Group differences were also observed in the EPM for open arm entries (*F*_(3,44)_ = 27.3, p < 0.0001, Figure 7c) and duration (*F*_(3,44)_ = 34, p < 0.0001, Figure 7c) and in the FST for immobility (*F*_(3,44)_ = 48.2, p < 0.0001) (Figure 7d). CORT+FLX non-responders that received DG Activin A had increased open arm entries and duration and decreased immobility relative to responders that received vehicle, Inhibin A, or Activin A+Inhibin infusions (p < 0.0001 for all, Bonferroni-corrected). Activin A+Inhibin coinfusions were not significantly different than vehicle or Inhibin A for EPM open arm entries, open arm duration, or FST immobility (p > 0.9999 for all, Bonferroni corrected). The negative affect index also demonstrated group differences (*F*_(3,44)_ = 538, p < 0.0001) (Figure 7e), with CORT+FLX non-responders that received DG Activin A infusions showing decreased negative affect relative to vehicle, Inhibin A, and Activin A+Inhibin infusions (p < 0.0001 for all, Bonferroni corrected). Significant relationships also emerged when we directly compared NSF latency to feed to EPM open arm duration (Pearson *r* = −0.821, p < 0.0001), with linear regression line (*y* = −5.23x − 826, *F*_(1,46)_ = 95, p < 0.0001), and to FST immobility (Pearson *r* = 0.863, p < 0.0001), with linear regression line (*y* = 3.67x − 139, *F*_(1,46)_ = 135, p < 0.0001) (Figure 7f). Taken together, these data demonstrate that coinfusion of Inhibin A blocks the ability of chronic DG Activin A infusions to convert FLX non-responders into responders across NSF, EPM, and FST, and further suggests that the effects of Activin A infusions are mediated through downstream Activin signaling.

**Figure 7.**
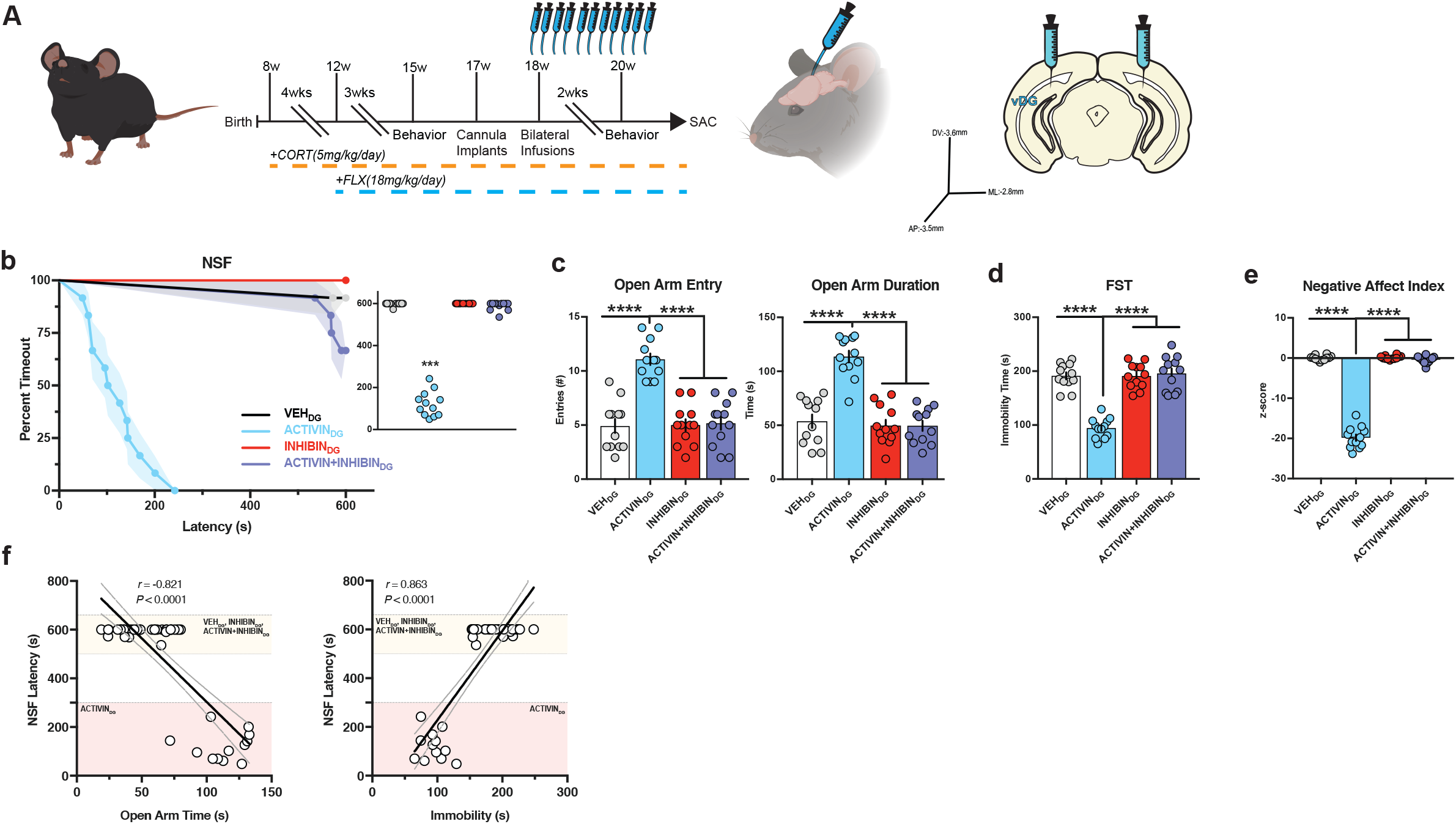
Coinfusion of Activin A and Inhibin A into Non-Responders blocks the effects of Activin A on behavior. (a) Timeline of experiment and coordinates of infusions for ventral DG. (b) Kaplan-Meier survival curve (large panel) and scatterplot (small panel) of NSF data showing individual latency to eat values across all four non-responder treatment groups. (c-e) One-Way ANOVA of VEH, Activin A infusions into DG (ACTIVINDG), Inhibin A infusions into DG (INHIBIN_DG_), and combined Activin A and Inhibin A infusions into DG (ACTIVIN+INHIBINDG) for EPM open arm entries (c left panel), EPM open arm duration (c right panel), FST immobility (d), and Negative Affect Index (e). (f) Regression analyses correlating NSF latency to eat with EPM open arm duration (left) and FST immobility (right). For survival curves, line shading shows SEM of each group (n=12 per group). Scatterplots, horizontal lines, and bars show group means with errors bars indicating SEM.

### Activin A infusions into DG are a more effective augmentation therapy than commonly used second-line treatments

When human patients do not remit to initial antidepressant therapy, they are usually switched to a new antidepressant. For example, in the large NIMH funded STAR*D study^2^, patients were first treated with citalopram (Celexa, a SSRI). Approximately 33% were found to display remission of depression symptoms. The 67% that did not remit were then subdivided into several groups and switched to either sertraline (Zoloft, a SSRI), bupropion (Wellbutrin, a norepinephrine/dopamine reuptake inhibitor), or venlafaxine (Effexor, a serotonin/norepinephrine reuptake inhibitor). Other groups either remained on citalopram and were augmented with bupropion or received other treatments. Our data suggests that chronic DG Activin A infusions are a very effective augmentation strategy for non-responders to FLX treatment. Therefore, we next wanted to assess whether switching mice from FLX to other antidepressants or augmenting FLX with other antidepressants is as effective in converting non-responders into responders as augmenting FLX with Activin A infusions into DG. To this end we exposed a large cohort (n=250) of C57BL/6J mice to chronic CORT+FLX and then assessed their behavior in NSF, where we found that n=79 were non-responders to FLX treatment. Two weeks later, we cannulated all 79 FLX non-responders to bilaterally target the DG and then housed the mice two per cage with a divider (timeline of experiment in Figure 8a). One week after cannulation, we subdivided these non-responders into 6 groups of mice (n=12-14 per group). Two groups remained on FLX, one group was switched to sertraline (SER, 10mg/kg/day)^42^, one group was switched to bupropion (BUP, 10mg/kg/day)^43^, one group was switched to venlafaxine (VEN, 20mg/kg/day)^44^, and the remaining group remained on FLX but also began receiving bupropion (FLX+BUP, 10mg/kg/day of BUP). Then, one week after the groups were formed, we began bilateral infusions. One of the two groups that remained on FLX alone received Activin A infusions, while the other five groups received vehicle infusions. Infusions were given once per day (over a time course of 15 minutes per hemisphere) for two weeks. We then retested these six groups of mice in NSF (Figure 8b). The mice that remained on FLX only and received vehicle infusions remained non-responders. Consistent with the results in Figures 5 and 7, 100% (12/12) of the mice that remained on FLX alone and received Activin A infusions into DG showed decreased latency to eat and were converted into responders (p < 0.0001 relative to FLX alone, logrank Mantel-Cox test with Bonferroni correction) (Figure 8b-c). By contrast, only 28.5% (4/14) of the mice switched to sertraline (p = 0.0408 relative to FLX alone, logrank Mantel-Cox test, not significant with Bonferroni correction), 30.8% (4/13) of the mice switched to bupropion (p = 0.0329, logrank Mantel-Cox test, not significant with Bonferroni correction), 35.6% (5/14) of the mice switched to venlafaxine (p = 0.0193, logrank Mantel-Cox test, not significant with Bonferroni correction), and 38.5% (5/13) of the mice that remained on FLX and were augmented with bupropion (p = 0.0145, logrank Mantel-Cox test, not significant with Bonferroni correction) were converted into responders. Therefore, only the chronic DG Activin A infused group showed a significantly decreased latency to eat in NSF relative to the FLX only group. These data strongly suggest that direct modulation of Activin signaling in the DG may be a more effective augmentation strategy than those that are currently used.

**Figure 8.**
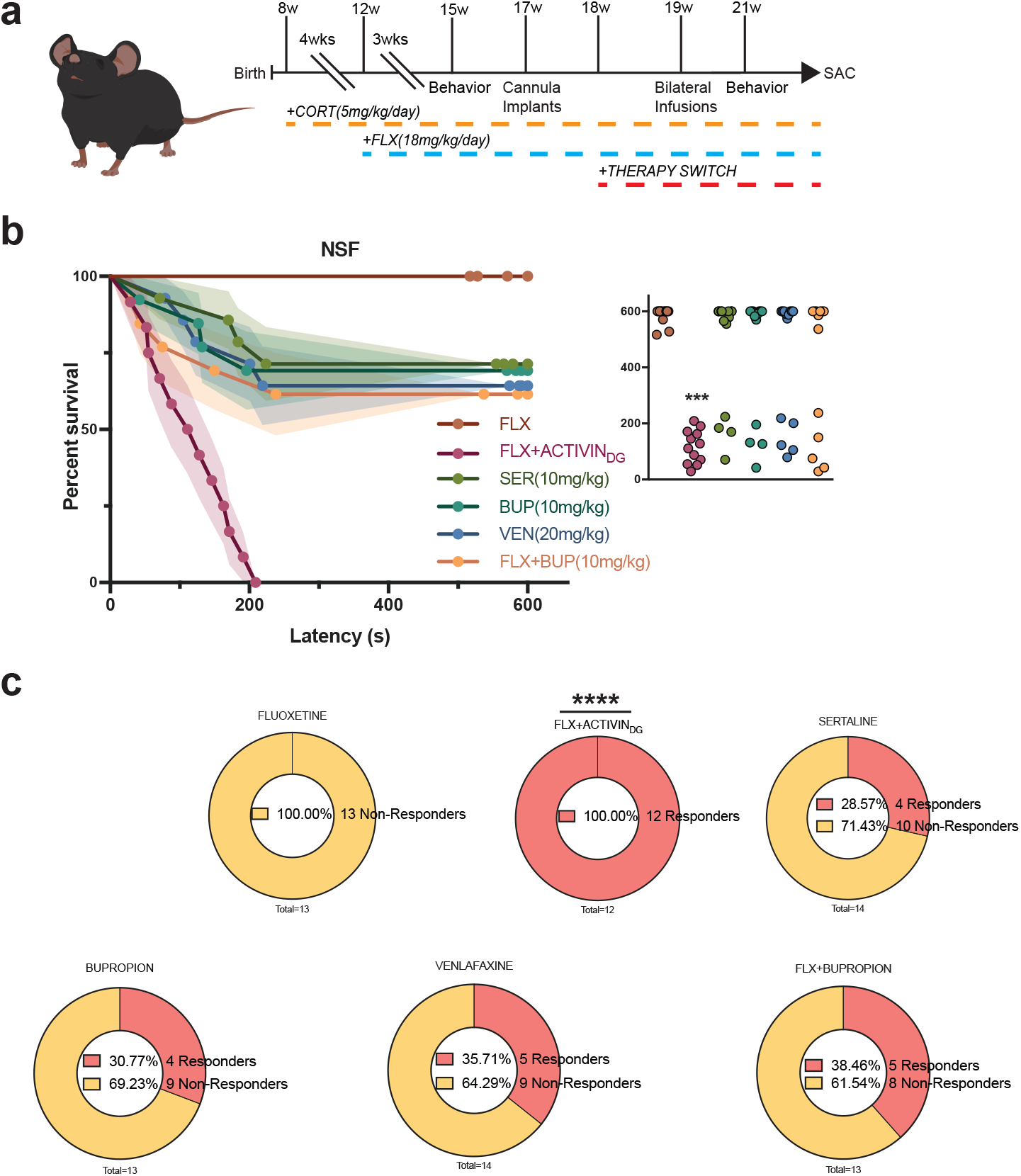
Activin A infusions into DG are a more effective augmentation therapy than commonly used second-line treatments. (a) Timeline of experiment. (b) Kaplan-Meier survival curve (large panel) and scatterplot (small panel) of NSF data showing individual latency to eat values across all six CORT+FLX non-responder treatment groups. (c) Graphical depiction of proportion of CORT+FLX non-responders converted into responders following the different second-line treatments.

## DISCUSSION

Our results demonstrate that FLX response status is conserved across several negative valence behavior tests (mice that show a response to FLX in NSF also show a response in EPM and FST and vice versa for non-responders) regardless of whether chronic corticosterone or chronic social defeat stress was used to induce a negative affective state. Furthermore, FLX response status was highly correlated with DG expression of multiple Activin signaling components. Functionally, chronic activation of Activin signaling in the DG successfully converted behavioral non-responders to FLX into responders. By contrast, chronic inhibition of Activin signaling in the DG converted responders to FLX into non-responders. This bidirectional modification is the first evidence that response or resistance to an antidepressant can be altered. Furthermore, these results strongly suggest that Activin signaling in the DG is a necessary component of achieving a behavioral response to antidepressant. Finally, chronic activation of Activin signaling proved to be a more effective augmentation strategy for non-responders to FLX than several commonly used second-line treatments.

### Antidepressant Treatment Resistance

Within the United States, approximately 16% of the population will experience an episode of major depression in their lifetime^1^. Although commonly used treatments, such as SSRIs, are prescribed to relieve symptoms, only a subset of patients (~33%) achieve remission with initial treatment^3^. Given that SSRIs are also prescribed widely for several anxiety disorders and obsessive-compulsive disorder, this treatment resistance results in clinicians using decision-tree medical algorithms, such as the Texas Medication Algorithm Project (TMAP)^45^, in attempts to combat mood disorders in patients that do not remit to initial lines of treatment. As patients move through different levels of these treatment algorithms, remission rates drop dramatically (~35% with second treatment to ~16% with fourth treatment)^2,46,47^. Therefore, failure to achieve remission within two treatments results in very poor outcomes. These issues plague modern psychiatry and indicate that drugs targeting monoaminergic systems have reached a limit in terms of effectiveness. Much research now focuses on developing drugs that take aim at distinct targets, including glutamate modulators, anticholinergic agents, and opioid modulators rather than monoaminergic systems^48–53^. However, it remains unclear why SSRIs and other monoaminergic drugs are only effective for a subset of patients. Our unique approach here to assess SSRI treatment resistance in mice suggests that individual molecular differences within the neural circuitry underlying the antidepressant response may underlie response status. Direct activation of Activin signaling in the DG proved to be more effective in converting non-responders to FLX treatment into responders. Therefore, similar directed molecular or even neural circuit-based approaches may ultimately prove to be more effective augmentation strategies than blindly switching treatments.

### Dentate Gyrus is a critical component of the neural circuitry mediating the antidepressant response

We assessed molecular differences between FLX responders and non-responders in the DG of mice exposed to chronic stress. Our data suggests that manipulation of DG Activin signaling can bidirectionally alter the behavioral response to FLX. These data further support a growing preclinical literature utilizing ablation, genetic, and neural circuit-based approaches to demonstrate that the DG is a principal component of the neural circuitry regulating both mood and the antidepressant treatment response^6,9,11,12,19–22^. It remains to be determined whether neural circuit-based approaches to modify the DG can alter antidepressant response status in a similar fashion to Activin A infusions.

Several distinct populations of cells in the DG, including the young adult born granule cells (younger than 8 weeks)^7,9,11^, mature granule cells (developmentally born or adult born granule cells older than 8 weeks)^6^, and cytocholecystokinin (CCK)-positive GABAergic interneurons^22^ are implicated in mediating the antidepressant treatment response. In all likelihood, these distinct populations work in concert via local microcircuitry. One common thread among these cell types is that the young adult born granule cells and CCK-positive interneurons provide an inhibitory influence over the mature granule cells in the ventral DG that is critical for both stress resilience and the antidepressant response^20,22^. Furthermore, inhibitory 5-HT1A receptors on mature granule cells are required for the antidepressant response^6^. Our preliminary microarray data that implicated Activin signaling components in the antidepressant response and all data in this manuscript were from microdissections of the granule cell layer (GCL) of the DG^38^, which is primarily composed of densely packed mature granule cells and sparse young adult born granule cells^54^. Therefore, we hypothesize that the Activin signaling components are being altered within these cell types. However, it will be important for future work to detail the exact effects of Activin signaling on DG neuronal ensembles, and how Activin signaling affects DG activity and ultimately other components of the neural circuitry underlying the antidepressant response.

### The role of Activin signaling in the antidepressant response

The importance of Activin and Inhibin signaling, as part of the TGF-β superfamily, is well-understood in the context of development, where Activin plays important roles in erythroid cell differentiation, induction of the dorsal mesoderm, and craniofacial development^41,55,56^. Activin also plays an essential role in pituitary follicle-stimulating hormone (FSH) production, while Inhibin inhibits FSH production^41,55^. However, the roles of these protein complexes are less understood in the context of the developed brain. While basal levels are low, Activin A is rapidly induced in the hippocampus by electroconvulsive seizures and long-term potentiation (LTP)-inducing high frequency stimulation, where it plays a role in the maintenance of long-term memory and late-LTP^57–60^. Activin A and Acvr1A mRNA are upregulated in the DG by chronic treatment with the antidepressant paroxetine and Smad2 phosphorylation is induced by fluoxetine treatment^16,17^. Environmental enrichment (EE) induces Activin A mRNA expression in DG and CA3 and increases Smad2/3 phosphorylation in the hippocampus^60^. Overexpression of a dominant-negative Acvr1B in mouse forebrain under the CamKIIa promoter resulted in anxiolytic-like behavioral effects, a decreased behavioral response to benzodiazepines, enhanced spontaneous GABA release, and increased GABA tonus^61^. Inducible transgenic expression of Activin A under the CamKIIa promoter resulted in anxiolytic-like behavioral effects in the Open Field, EPM, and Light-Dark Test, while expression of Follistatin, an inhibitor of Activin signaling, under the CamKIIa promoter had anxiogenic effects^62^. Furthermore, acute Activin A infusions into DG but not CA1 or Amygdala, reduces immobility in FST^16,17^. Acute Activin B infusions had no effect. Finally, a human genetic association study found 166 single nucleotide polymorphisms (SNPs) within 10 genes belonging to the Activin signaling pathway as being associated with antidepressant treatment response^17^. Genetic variants in the betaglycan gene *(TGFBR3),* a member of the human Activin system, showed the best association, as homozygote carriers of the major allele were significantly more frequent among the responders to antidepressant treatment^17^. Taken together with our results, it is now clear that Activin signaling in the DG is a critical component of the behavioral response to antidepressant treatment.

## ACKNOWLEDGEMENTS

This work was supported by: NIMH (R01MH112861 and K01MH098188, B.A.S.) and BBRF NARSAD Young Investigator Award (B.A.S.). The authors thank Dr. Rene Hen for his guidance and mentorship throughout this project, Alice Hu for technical support, and Drs. Dirk Moore and John McGann for their advice on statistical analyses.

## AUTHOUR CONTRIBUTIONS

B.A.S. conceived experiments. B.A.S., M.R.L., C.N.Y., and M.M.G. performed the experiments. B.A.S., M.M.G., and C.N.Y. analyzed the data and made the figures. C.N.Y., M.M.G., and B.A.S. wrote the manuscript.

## METHODS

### Mice

Adult 8-week-old male mice purchased from Jackson labs were housed in groups of three to five per cage with *ad libitum* access to food and water. Mice were on a 12:12-h light/dark schedule. All behavioral testing was conducted during the light period and using the same testing time throughout the experiments. Mouse protocols were approved by the Institutional Animal Care and Use Committee at either Columbia University or at Rutgers, The State University of New Jersey.

### Drug administration

Animals were either placed on chronic doses of either vehicle, which consisted of 0.45% β-cyclodextrin in water, or corticosterone (5mg/kg) dissolved in vehicle for duration of experiments. After 4 weeks of either vehicle or corticosterone administration, a subgroup of animals received either vehicle (autoclaved water) or fluoxetine (18mg/kg) via daily oral gavage for throughout the experiment. On the days when mice were subjected to behavioral testing, fluoxetine or vehicle administrations were conducted after the mice completed the testing in order to avoid any acute effects. For additional therapies mice were then treated for 3 weeks with either: bupropion (10mg/kg), venlafaxine (20mg/kg), sertraline (10mg/kg), or fluoxetine + bupropion (18mg/kg, 10mg/kg, respectively) via oral gavage.

### Behavioral testing

Behavioral testing was conducted after 3 weeks of antidepressant administration, in the following order: EPM, NSF and then FST. Mice were given 3 days between behavioral tests to avoid contaminating stressors as well as before sacrifice. Prior to each behavioral test mice were acclimated to the room for 30 mins of habituation.

#### Elevated plus maze

EPM was performed as previously described^6^. The plus maze consisted of two closed arms and two open arms 2 feet above the floor. Mice were placed into the central area facing one closed arm and allowed to explore the maze for 5 mins. Data were scored using Ethovision software (Noldus), in which open arm time and open arm entries were recorded. Between animals, the maze was cleaned with 70% ethanol between each run.

#### Novelty-suppressed feeding

NSF was performed as described^6,63^. The testing apparatus consisted of a plastic box (50 × 50 × 20 cm), the floor of which was covered with approximately 2 cm of bedding. 18 h before behavioral testing, mice were weighed and food deprived. At the time of testing, a single pellet of food was placed on a white paper platform in the center of the box beneath a gooseneck lamp illuminating the center of the area at about 1500 lux. A mouse was placed in a corner of the box and latency to approach and eat the food pellet was recorded with a maximum of 10 mins. Immediately after the testing period, the mice were transferred to their home cages and latency to feed in the home cage as well as consumption was recorded. After completing testing, animals were weighed again and % change in body weight was calculated with respective to pre-testing weights.

#### Forced swim test

As previously described^6^, mice were placed into clear plastic buckets 20 cm in diameter and 23 cm deep, filled two-thirds of the way up with 26 °C water and were videotaped. Mice were in the forced swim buckets for 6 mins, but only the last 4 mins were scored. Scoring was automated using Videotrack software (ViewPoint).

### Social Defeat Stress

Paradigm was conducted as previously described. In brief, defeat stress was carried out using similar methods to those already published^20,31^. Prior to social defeat stress, male retired breeder CD1 mice (Charles River Labs) were screened for aggression. CD1 mice that attacked screener mice (C57/BL6J) for two consecutive days in less than 60 seconds were selected. After selection of aggressors, experimental mice were exposed to a different CD1 aggressor mouse each day for 5 min over 10 days. After contact, experimental mice were inspected for injury and separated from the aggressor and placed in an adjacent compartment of the same cage as the CD1 mouse, separated by a plastic divider with holes. Control test mice were housed in equivalent cages but with members of the same strain, which were changed daily. Twenty-four hours after the last session, all mice were housed individually for the remainder of the study. Following 10 days of social defeat stress, animals were run through a social interaction test in which mice were placed in an open field for 2.5 minutes for baseline exploration in the absence of a novel CD1, and then for another 2.5 minutes in the presence of a CD1. Mice were deemed susceptible if they spent less time in the interaction zone when the social target was present than absent AND a total time spent interacting with the social target <30s, and resilient if the they spent more time in the interaction zone when the social target was present than absent AND total time spent interacting with the social target >60s.

### Gene Expression

Animals were sacrificed via rapid decapitation and dentate gyrus was microdissected, flash frozen, and then stored at −80°C until further processing. RNA was extracted from tissue samples using a RNA/DNA Purification kit (Norgen Biotek). Total RNA was then converted into cDNA using Superscript III enzyme (Invitrogen). Quantitative-PCR was performed in triplicate reactions with Taqman Fast Advanced Mastermix and Taqman probes for activin a, acvr1a, acvr1b, acvr1c, smad2, smad3, and rn18s (Life Technologies) on a StepOne Plus Real-Time PCR System (Applied Biosystems). Data was analyzed using the ΔΔC_T_ method, triplicate cycle thresholds per gene per sample were averaged, normalized to control gene *(rn18s)* to obtain ΔC_T_, and were then converted to ΔΔC_T_ values by normalizing to mean ΔC_T_’s of the vehicle group. Final values were then expressed as an expression percentage relative to the vehicle group values.

### Intracerebral infusions

Mice were anesthetized with sodium pentobarbital (diluted 1:10 from stock of 50mg/ml and injected at a volume of 10 ml/kg) and guide cannulae with dummy cannulae (Plastics 1) were implanted. For ventral DG the coordinates used were: −3.5 mm and ±2.8 mm from bregma at a depth of 3.6mm from the skull surface, and for CA1 the coordinates used were: −3.1mm and ±3.0mm from the bregma at a depth of 2.0mm from the skull surface. 1-2 weeks after surgery, animals began to receive bilateral infusions of either vehicle (0.1% BSA), 1μg of mouse Activin A peptide (R&D Systems), 1μg of mouse Inhibin A peptide (R&D Systems), or 1μg each of Activin A+Inhibin A in vehicle once per day (over a time course of 15 minutes per side, 10 minutes of infusion and an additional 5 minutes with tubing left in place) for 2 weeks prior to behavioral testing. Each day connector assemblies with tubing were connected to internal cannulae, which were then inserted into the guide cannulae. Infusions were delivered by a standard infusion only syringe pump (Harvard Apparatus). A total volume of 1.0 μl was infused in each hemisphere per day. Animals were freely moving in their cage during infusions.

### Statistics

All statistics were performed using Prism Software 7 (Graphpad). Parametric hypotheses were assessed with parametric tests. Two-way ANOVAs assessing stress pretreatment x antidepressant treatment were used for EPM, FST, Negative Affect Index, and gene expression. One-way ANOVAs were then subsequently used to assess Stress+VEH, Stress+FLX responders, and Stress+FLX non-responders. NSF was assessed using non-parametric Kaplan Meier survival analysis with log-rank Mantel-Cox. Correlational analysis between individual animal behavioral values (NSF and either EPM or FST) were performed using Pearson *r* and best-fit values with a linear regression analysis and slopes with analysis significant non-zeroes were analyzed. Post hoc Bonferroni corrections were used where appropriate.

---

**Supplemental Figure 1.**
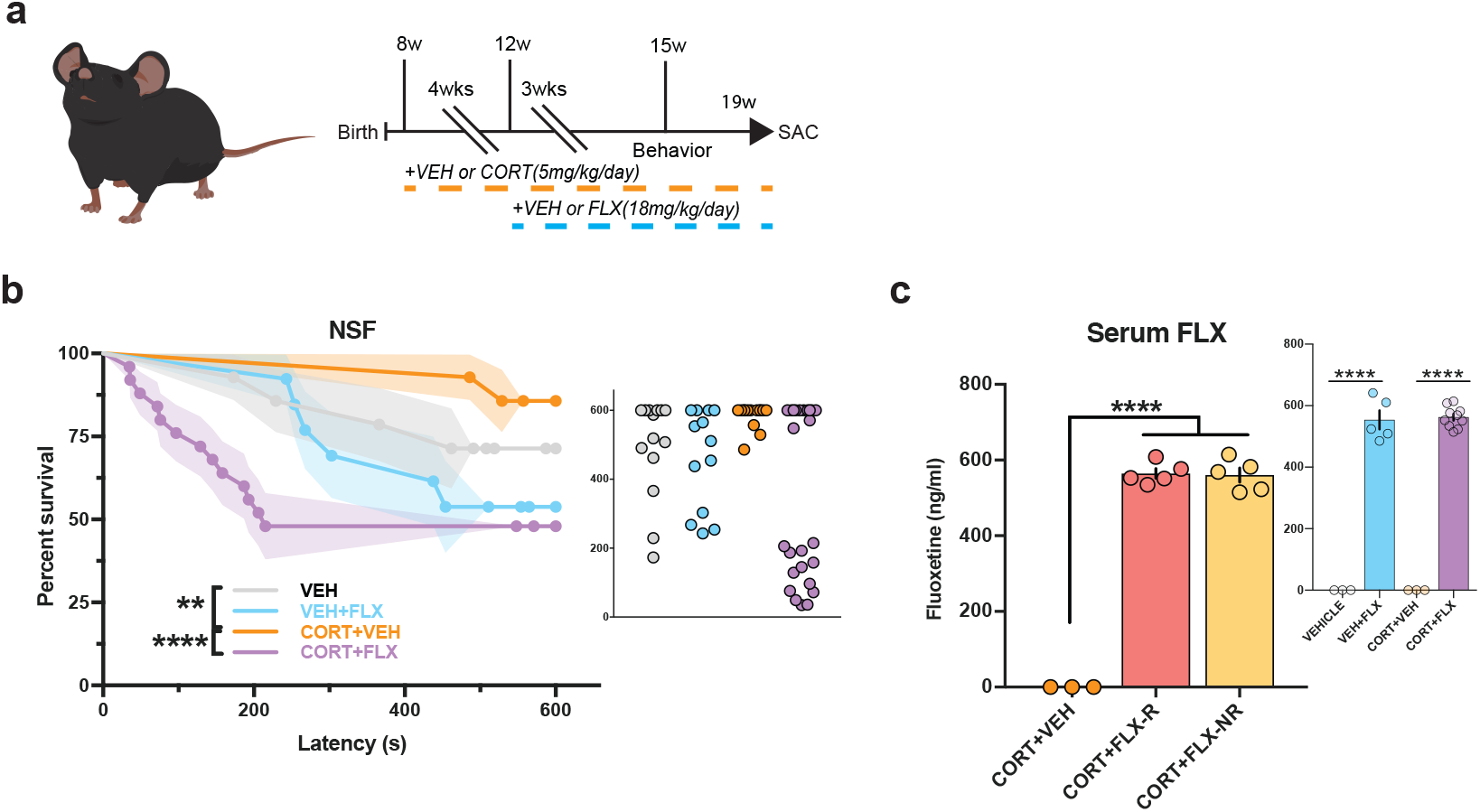
NSF data for DG mRNA expression cohort of mice and serum FLX levels. (a) Timeline of experiment. (b) Kaplan-Meier survival curve (large panel) and scatterplot (small panel) of NSF data showing individual latency to eat values across all four treatment groups. (c) Two-way ANOVA of all treatment groups (small panel) and One-Way ANOVA of CORT+VEH, CORT+FLX responders and CORT+FLX non-responders (large panel) for serum FLX levels following three weeks of FLX administration. For survival curves, line shading shows SEM of each group (n=15-23 per group). Scatterplots, horizontal lines, and bars show group means with errors bars indicating SEM.

**Supplemental Figure 2.**
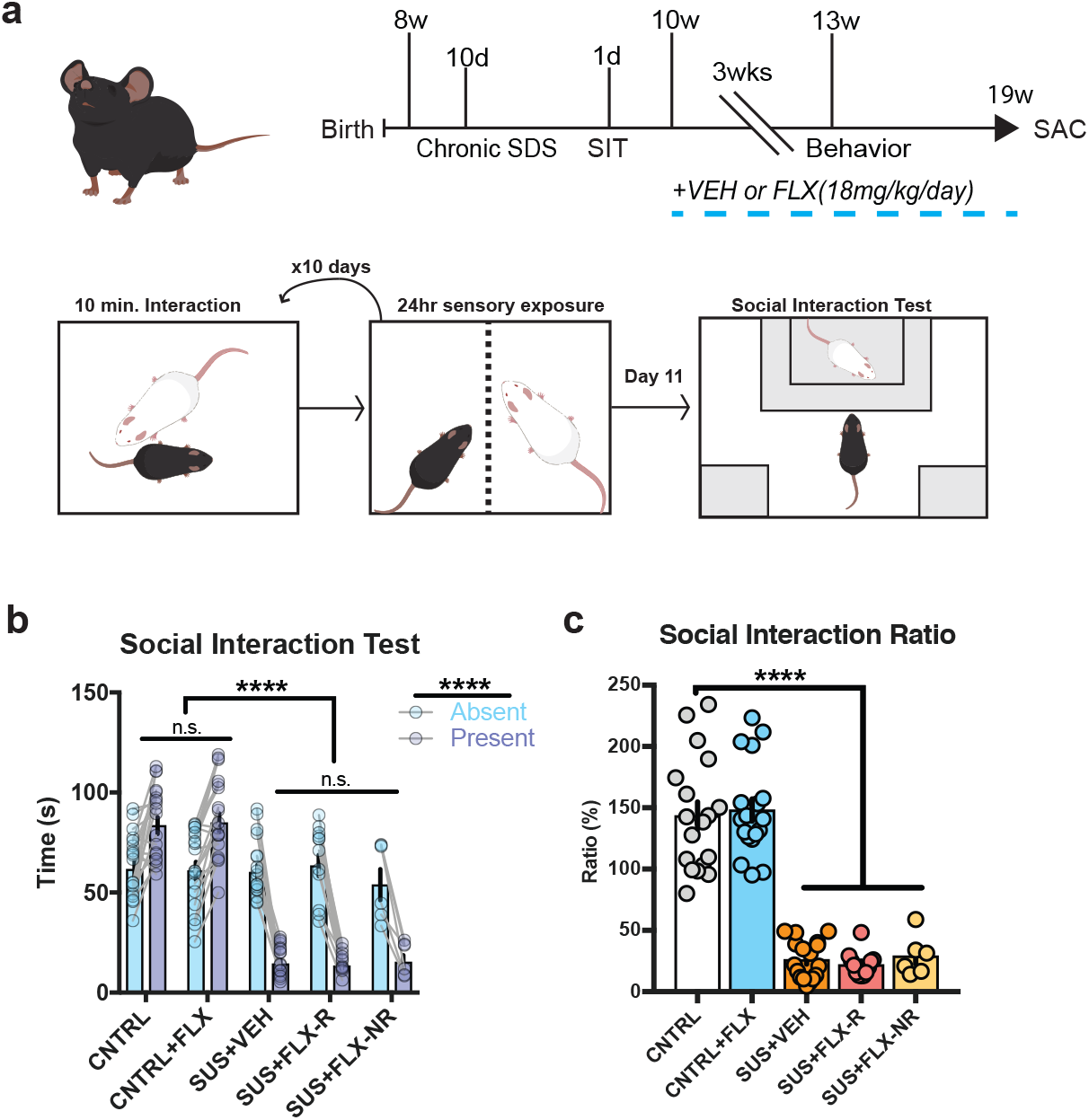
Identification and classification of susceptible mice in Figures 3 and 4. (a) Timeline of experiment and diagram of CSDS and Social Interaction (SIT) protocol. (b) Time spent in social interaction zone reveal significant difference (*F*_(3,67)_ = 106, p < 0.0001) only in the presence of a CD1 aggressor. The only difference between groups was CNTRL animals compared to SUS+VEH (p < 0.0001, Bonferroni-corrected), SUS+FLX-R (p < 0.0001, Bonferroni-corrected), or SUS+FLX-NR (p < 0.0001, Bonferroni-corrected). (b) Social interaction ratios were significantly different (*F*_(4,66)_ = 62.2, p < 0.0001). The only difference between groups was CNTRL animals compared to SUS+VEH (p < 0.0001, Bonferroni-corrected), SUS+FLX-R (p < 0.0001, Bonferroni-corrected), and SUS+FLX-NR (p < 0.0001, B onferroni-corrected).

